# The Expression of Angiopoietin-1 and −2 in the Osteogenesis of Mesenchymal Stem Cells

**DOI:** 10.1101/2021.04.15.440087

**Authors:** Jinwen Chen, Guangchan Yang, Jie Guo, Yuqin Liu, Jinchen Guo, Jiatao Suo, Hongyou Yu

**Affiliations:** Undergraduate Student, School of Medicine, Dalian University, P.R.China; Postgraduate student, School of Medicine, Dalian University, P.R.China; Department of Orthodontics, School of Medicine, Dalian University, P.R.China

**Author notes:** **Corresponding author**: Dr. Hongyou Yu.

**Keywords:** Angiopoietin-1, Angiopoietin-2, Mesenchymal Stem Cells

## Abstract

**Objective:** The objectives of this study are to clarify whether rat bone marrow derived Mesenchymal stem cells (MSCs) express Ang1 and Ang2 and their expression in the process of osteogenesis in vitro.

**Material and Methods:** MSCs were cultured from rat tibia bone marrow cells and the hemopoietic stem cells were deplete by consistently replacement of the culture medium. The MSCs were induced osteogenesis with mineralization conditional medium and Immunohistochemical and immunofluorescent staining were performed to assess the expression of Ang1 and Ang2.

**Results:** The method used to expand rat MSCs in vitro was applicable, and the cell morphology is spindle-like shape that is consistent with the privous reports. The immunohistochemical staining results showed that both Ang1 and Ang2 were expressed by rat MSCs. Both Ang1 and Ang2 were up-regulated in the process of osteogenesis of rat MSCs.

**Conclusion:** Rat MSCs express both Ang1 and Ang2 which might play critical roles in the osteogenesis in vitro.

## Introduction

Angiopoietins (Ang) are a family of cytokines that are associate with angiogenesis and inflammation. Human Angs include four members namely Ang1-4 while only Ang1 and Ang2 have been intensively investigated recently [Cascone and Heymach, 2012]. Ang1/2 is expressed in a variety of human tissues and the functions of Ang are related to angiogenesis and inflammatory response [2-3]. Both Ang1/2 bind to the receptor Tie2. Ang1 is an agonist of Tie2 which induces the Tie2 phosphorylation and stabilizes the vasculature while Ang2 is an antagonist of Tie2 which destabilizes the vasculater to initiate the angiogenesis. Ang-2 plays an important role in angiogenesis and inflammatory response, and is closely related to angiogenesis under physiological and pathological conditions [Fujiyama et al., 2001]. In the research of tumor [De Palma and Naldini, 2011], it is found that Ang is closely related to tumor angiogenesis, invasion and metastasis, and prognosis.

Bone marrow derived mesenchymal stem cells (Mesenchymal Stem Cells, MSCs) are stem cells with plural differentiation ability, which can differentiate into adipocytes and osteoblasts in vitro [Abdallah et al., 2019]. MSC can induce differentiation into osteoblasts in vitro, and is currently considered to be potentially applied for the treatment of mineralized tissue related diseases. Bone remodeling is the balance between the osteoblastic function of bone formation and resorption and accompanied by the coupling effect of neovascularization and inflammatory response [Bhattarai et al., 2016]. As Angs have played vital roles in the formation of new blood vessels and in inflammatory response [Imhof and Aurrand-Lions, 2006], diabetic wound healing and cardiovascular disease [Harfouche and Hussain, 2006, Hozzein et al., 2018], this research would explore the expression of Ang1/2 in the process of MSC differentiation into osteoblasts, and provide theoretical basis mechanisms for the application of Angs in the process of bone remodeling, which is great significance for explore new strategies for clinical treatment of mineralization dysfunction related diseases, such as osteoporosis.

Current studies have shown that Ang is expressed in adipocytes, but there is still no direct evidence for the expression of Ang in MSCs [Kim et al., 2013], and there are few reports on the expression of Ang in the osteogenesis of MSCs in vitro. Therefore, this research has investigate the expression Ang1/2 during the osteogenesis of MSCs in vitro.

## Materials and methods

### Mesenchymal Stem Cell Culture

The protocol was performed following the guidelines approved by the Institutional Animal Care and Use Committee of Dalian University. The 3 week-old rats were euthanized and the tibias were dissected and were cut open on both ends. A 1ml syringe was used to flush the bone marrow cells out with 1 ml PBS until the tibia showed paled. The flushed cell were centrifuged at 1000rpm for 5 mins and then resuspended with DMEM medium(Solarbio) supplemented with 20% heat-inactivated fetal bovine serum (FBS, Solarbio) and 1% penicillin/streptomycin(Solarbio). The suspended cells were cultured in a 25cm cell culture flask (Nest) in the incubator with a humidified atmosphere 5% CO_2_ and 37°C. The medium was initially replaced in 6 hours followed with every 24 hours for 3 days. The attached fibro-liked cells cells wer MSCs. When the cells grew more than 80% confluence, 0.25% trypsin EDTA was applied to detached the cells that were passed to expand the cells. The passge 3-6 were used for this experiment.

### Osteogenic differentiation and mineralization node staining

MSCs were induced osteogenesis by supplementing cells with 100 nM dexamethasone, 10 μM β-glycero-phosphate, and 50mg/ml L-acid ascorbate (all from Sigma-Aldrich) for 2 weeks. The cells were fixed at day 7 and 14 that are used for the following immunohistochemical and immunofluorescent staining.

### Immunohistochemical staining

The attached cells were harvested and washed with cold PBS for three times followed with a fixation using 100% ice cold methanal at −20°C for immunohistochemical staining. Briefly, the cells were then incubated with 10% goat serum (Solarbio) for 30 mins at room temperature. Then the cells were incubated with 1:1000 rabbit anti-Ang1 (Proteintech) or 1:1000 mouse anti-Ang2 (Santa Cruz) for 1 hour at room temperature. After washing with PBS for 3 times, the cells were incubated with 1:5000 HRP conjugated goat anti-rabbit IgG or goat anti-mouse IgG (both from Solarbio) for 1 hour at room temperature. The cells were then washed three times with PBS and DAB and hematoxylin counterstaining were performed as per manufacture protocol (both from Solarbio). Images (1600 x 1200 pixels) were acquired at 40x magnification at standardized settings using a Olympus microscope.

### Immunofluorescent staining

The immunofluorescent staining was similar as the immunohistochemical staining above. The secondary antibody 1:5000 Coralight 488 goat anti-Rabbit (Proteintech) and 1:5000 coralight 584 goat anti-mouse (Proteintech) was applied for 1 hour at room temperature in dark. DAPI was used to counterstaining the nuclear (Solarbio) for 5min. Images (1600 x 1200 pixels) were acquired at 40x magnification at standardized settings using an Olympus microscope. ImageJ (NIH) software was used to analysis the average fluorescence intensity following the instruction by NIH ImageJ software.

### Statistical Analysis

The statistical analysis were performed with the free statistical software GNU PSPP. Two-sample *t*-test was used to analysis the expression for Ang1 and Ang2 between groups with and without osteogenesis of MSCs at different time points. Data are presented as mean ± SD and P<0.05 was considered statistical significant.

## Results

### Morphological characterization of rat Mesenchymal stem cells

Primary cell culture was initiated after constantly medium replacement at the first three days which could deplete the homeopathic stem cell contamination. The MSCs were observed and photographed under a phase-contrast microscope. The MSCs showed colony growth (Figure 1A-C) and single cells with spindle-shaped or vortex-shaped cells develops within 5d (Figure 1D). The cells reached about ∼90% confluence on day 8. The cell populations were morphologically homogenous fibroblast-like cells in the following passage.

**Figure 1.**
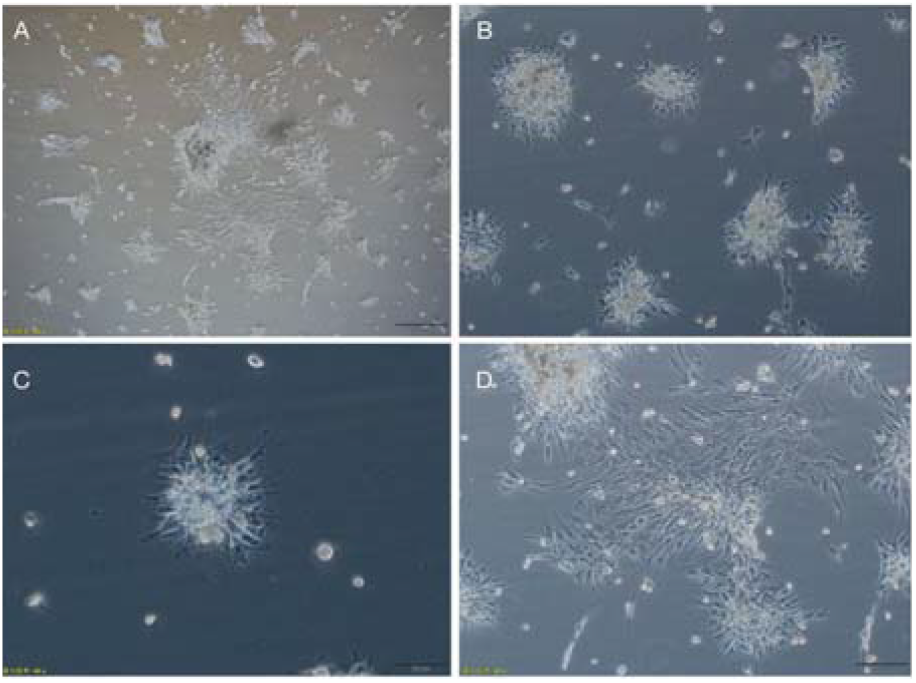
The morphology of the rat MSCs under the phage-contrast microscope. A. colonies formed at day 5, the magnification 20x; B. cell populations were observed under 40x magnification; C. single colony of the cells with a 40x magnification; D. cell morphology at day 8 with a 40x magnification. The scale bar length is 40μm(20x) and 20μm (40x)

### Rat MSCs express both Ang1 and Ang2

Immunohistochemical staining of Ang1 and Ang2 results showed that rat MSC expressed both Ang1 and Ang2 (Figure 2). The Ang1 expression is strongly expressed mainly around the nuclei (Figure2A) while the Ang2 expression is slightly less compared with Ang1 and it is evenly distributed in the cytoplasm (Figure 2B). These results indicated that rat MSCs express both Ang1 and Ang2 in cytoplasm.

**Figure 2.**
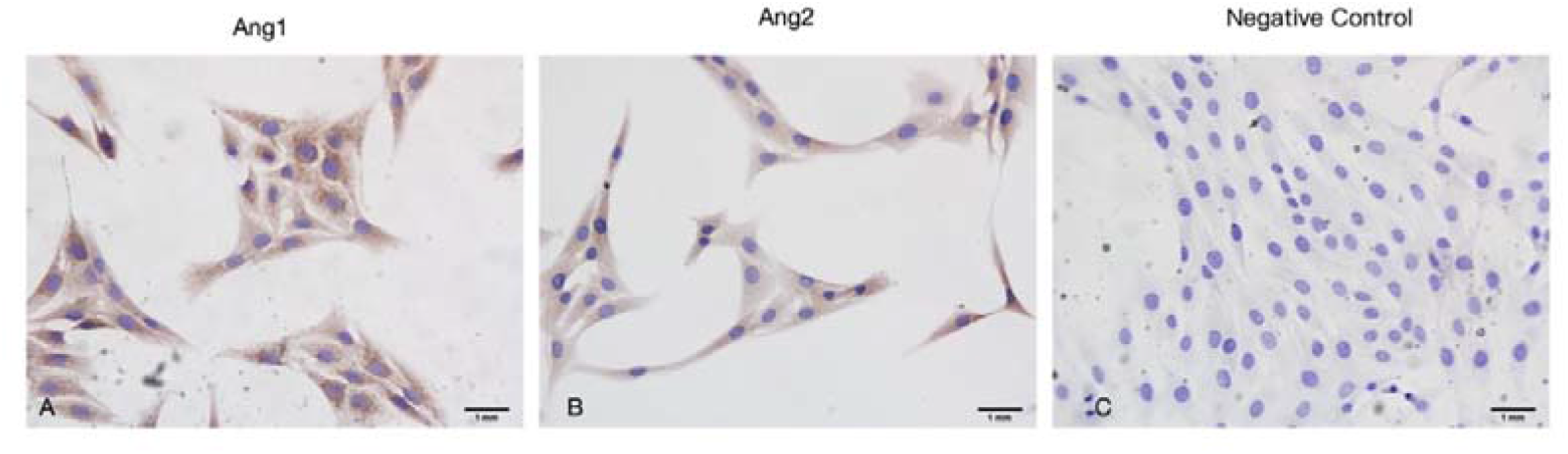
The expression of Ang1 and Ang2 on the rat MSCs. A. IHC staining of Ang1; B. IHC staining of Ang2; C. Negative control; the magnification 20x and the scale bar length is 1mm.

### The up-regulation of Ang1 and Ang2 expressing during osteogenesis of MSCs

Immunofluorescent staining results showed that rat MSCs expressed both Ang1 and Ang2 which is consistent with the results of immunohistochemical staining (Figure 3). During the process of osteogenesis, the Ang1 and Ang2 expression were both up-regulated (Figure 3).

**Figure 3.**
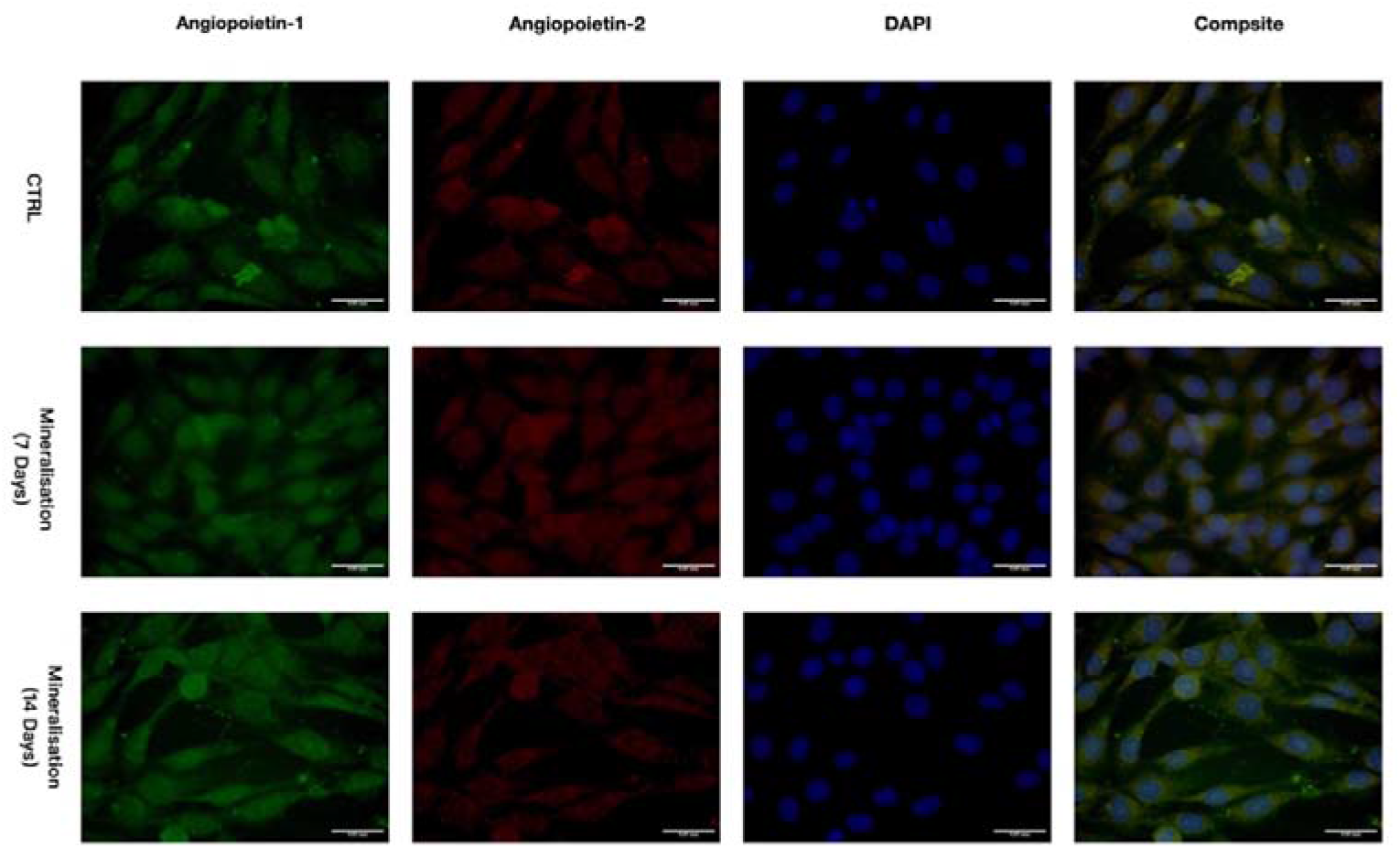
The represent images of the expression of Ang1 and Ang2 during the osteogenesis of MSCs. A. the expression of Ang1 and Ang2 in the rat MSCs; B. IHC staining of Ang2; C. Negative control; the magnification 20x and the scale bar length is 1mm.

Quantification the average immunofluorescent density of Ang1 and Ang2 on day 7 and day 14 osteogeneses showed that both of them were significant up-regulated (Figure 4). The average fluorescent intensity of Ang1 on day 7 and day 14 were statistically significant (186±8.89 and 222±10.21 on day 7 and 14 respectively vs 165±8.73 on the control group). The average fluorescent intensity of Ang2 on day 7 and day 14 showed similar trends as Ang1 (137±7.50 and 166±1.20 on day 7 and 14 respectively vs 118±8.08 on the control group).

**Figure 4.**
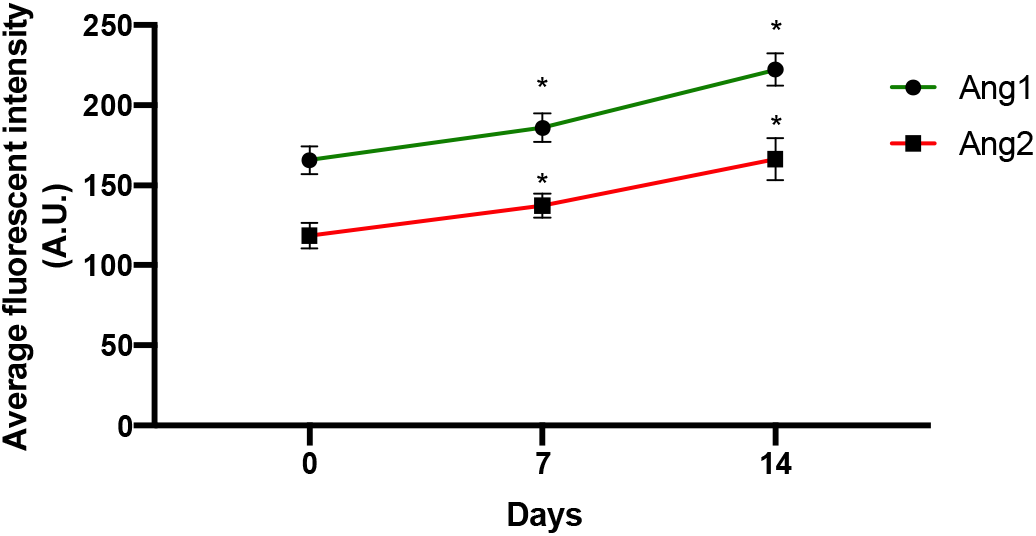
The average fluorescent intensity of Ang1 and Ang2 during osteogenesis of rat MSCs. *P<0.05 and the unit is Arbiter Unit (A.U.).

## Discussion

In this study, we investigated the expression of Ang1 and Ang2 in the MSCs and their expression in the process of osteogenesis in vitro. It was found that rat MSCs expresses both Ang1 and Ang2 and the expression of Ang1 and Ang2 are both up-regulated in the process of osteogenesis. The up-regulation of Ang1 is much stronger compared with Ang2.

Angs are mainly expressed by vascular endothelial cells, however, it has been found that Angs are expressed by various types of cellssuch as macrophages and osteoblasts. The functions of Angs are not only to induce angiogenesis and stablise the vasculature, it also has some important roles in regulation of inflammation [Imhof and Aurrand-Lions, 2006]. It has been found that the expression of Angs in osteoblast to niche the hompoietic stem cells in the bone marrow [Arai et al., 2004] and Ang2 plays critical roles in promoting the differentiation of bone marrow stem cells [Kang et al., 2017]and comp-ang1 promoting condrogenic and osteogenic differentiation of MSCs[Kim et al., 2013]. To our best knowledge, we have firstly investigated the expression of Ang1 and Ang2 in the rat MSCs and our results are consistent with previously study that the Ang1 and Ang2 were up-regulated in the process of osteogenesis of MSCs[Kim et al., 2013].

Bone marrow derived MSCs have the ability to induce plural differentiation of osteoblasts, chondrocytes, adipocytes and other cells under conditional medium in vitro. Besides it is minimal immune features, the plural differentiation ability makes MSCs suitable for the tissue regeneration for tissue repair, such bone defect repair [Bianco, 2014]. Bone, as a highly vascularized tissue, the growth efficiency and range of its blood vessels determines the rate of formation of new bone and the efficiency of bone defect repair [Dhillon et al., 2013], so it is necessary to make the artificial bone implanted in the body survive and play its due role. Promoting vascularization during new bone formation is the prerequisite and basic condition as it could not only delivery the bone precursor cells migration to the local tissue but also delivery the essential cytokines and nutrition that are essential for bone formation [Kasama et al., 2007]. Since both Ang1 and Ang2 plays critical roles in the osteogenesis of MSCs and Ang1 and Ang2 have critical roles in the angiogenesis. So it is feasible to proposed that the Ang1 and Ang2 might be best growth cytokines that could greatly facilitate the bone tissue repair via its ability to induce stem cells osteogenesis and new vessel formation. Indeed, Ying et al. [Yin et al., 2018] established animal experimental models to study Ang2 The mechanism that regulates autophagy and promotes the vascularization of tissue engineering artificial bone. Bhattarai et al. [Bhattarai et al., 2016] also early proposed the construction of tissue engineering bone for bone defect repair in the damaged mandible. In addition, Chen et al. [Chen et al., 2019] studied the effects of BMSCs transplantation on neurological function and the expression of Ang-1, Ang-2 and Tie-2 during the recovery period of cerebral ischemia in rats, and the results showed that transplantation of BMSCs can affect the neurological function of rats. When it is significantly improved, the signal pathways of Ang-1 and Tie-2 can produce endogenous new neurons and other effects. Xu et al. [Florian et al., 2021] showed that the bone marrow mesenchymal stem cells with high expression of the target gene Ang-1 constructed by genetic technology can detect the transformation and repair of MSCs in the lung after transplantation, which is a good gene carrier for the treatment of lung disease.

Bone defects have always been a problem in the fields of orthopedics, stomatology, and neurosurgery. MSC has been used for the treatment of many diseases, such as femoral head necrosis, bone regeneration, neurodegeneration, graft-versus-host, etc. Our study suggested that Ang1 and Ang2 might facilitate the bone formation. Therefore, it is applicable to combine MSCs and Ang1 and (or) Ang2 for clinical bone defect related tissue repair. However, the mechanisms of Ang1 and Ang2 expression regulation and the functions in the process of osteogenesis need future investigation.

## Conclusion

In this study, we confirmed that rat bone marrow derived MSCs express both Ang1 and Ang2 which are both up-regulated in the process of osteogenesis of MSCs.

## Acknowledgements

The simulations were performed on resources provided by the Swedish National Infrastructure for Computing at PDC (Center for Parallel Computing). We acknowledge the use of Fenix Infrastructure resources, which are partially funded from the European Union’s Horizon 2020 research and innovation programme through the ICEI project under the grant agreement No. 800858. The authors wish to thank Johanna Frost-Nylén, Robert Lindroos and Ilaria Carannante for helpful discussions. We also thank Robin de Schepper, Kadri Pajo, and Wilhelm Thunberg for help with software compatibility.

